# Aerobiology in High Latitudes: Evidence of Bacteria Acting as Tracer of Warm Air Mass Advection reaching Northern Antarctic Peninsula

**DOI:** 10.1101/2021.06.27.450079

**Authors:** Marcio Cataldo, Heitor Evangelista, José Augusto Adler Pereira, Álvaro Luiz Bertho, Vivian Pellizari, Emanuele Kuhn, Marcelo Sampaio, Kenya Dias da Cunha, Alexandre dos Santos Alencar, Cesar Amaral

## Abstract

Despite the extent use of geochemical tracers to track warm air mass origin reaching the Antarctic continent, we present here evidences that microorganisms being transported by the atmosphere and deposited in fresh snow layers of Antarctic ice sheets do act as tracers of air mass advection from the Southern Patagonia region to Northern Antarctic Peninsula. We combined atmospheric circulation data with microorganism content in snow/firn samples collected in two sites of the Antarctic Peninsula (King George Island/Wanda glacier and Detroit Plateau) by using flow cytometer quantification. In addition, we cultivated, isolated and submitted samples to molecular sequencing to precise species classification. Viable gram-positive bacteria were found and recovered in different snow/firn layers samples, among dead and living cells, their number concentration was compared to northern wind component, stable isotopes of oxygen, δ^18^O, and the concentration of crustal elements (Fe, Ti and Ca). Use of satellite images combined with air mass back-trajectory analysis obtained from the NOAA/ HYSPLIT model corroborated the results.

## 1. INTRODUCTION

The aerial transport/dispersal of biological material is believed to be an important component of the input to remote locations such as the Antarctic region, and plays a crucial role in shaping patterns of biodiversity in such places (Fierer, 2008; Pearce et al., 2010; Pearce et al. 2016). Intimately linked with ecological and evolutionary processes in the colonized environments, the rates of airborne input, survival after transport process, and viability after arrival, is essential for understanding the origin and dynamics of several processes. The capacity of cells to live in extreme environments and remain viable for thousands and millions of years (Priscu et al. 1998; Karl et al. 1999; Christner et al. 2003; Bidle et al. 2007) has attracted the attention of the scientific community to microorganisms found in deep and old ice of the Earth’s polar regions. Although the interest in the microbial communities of Antarctica has typically been turned toward ecologic, phylogenetic, and metabolic studies, microorganisms entrapped in snow/firn/ice layers in ice sheets could be looked from a different point of view, such as their origin and moment of deposition.

Considering the atmospheric transport events of short duration (time scale of days) between South America and the Antarctic Peninsula, the use of gaseous geochemical air mass tracers of crustal sources, for instance the ^222^Rn, have been successfully used, since the 80’s decade (Pereira, 1999). Furthermore, combined ^222^Rn and Si, anthropogenic elements (Bi, Cd, Cr, Cu, Ni, V, and Zn) and biomass burning aerosols (black carbon) were also employed as tracers (Evangelista and Pereira 2002; Mishra et al. 2004; Pereira et al. 2006). Like the geochemical atmospheric tracers, the presence of South-American plants pollen was also observed in the seas of the Antarctic region since the early decades of the 20^th^ century as well as a continuous arrival of pollen, by atmospheric transport, was firstly reported (Fritsch 1912 cited in Marshall 1996a).

The concept of monitoring biological material as atmospheric tracers is broader than the use of pollen species, since the Antarctic ice sheets contain a great diversity of microorganisms as viable and non-viable cells (Abysov 1993), and potentially they may indicate exogenous origin. It has earlier shown that these microorganisms are found in several different Antarctic microhabitats and many of these species are not able to duplicate themselves in a normal life cycle (Warwick 2000). This finding suggests that these species may arrive in Antarctica coming from other regions of the globe, and may not derive from local communities (Warwick 2000). Additionally, the Antarctic habitats support many microorganism species of cosmopolitan nature (Warwick 1988). Therefore, the Antarctic continent, besides its geographic and climatic isolation, seams to continuously exchange biological matter with other regions of the planet (Warwick 1988, 2000). Microorganisms are transported to continental scales mostly attached to dust or organic fragments (Prospero et al. 2005), but till now their atmospheric monitoring in Antarctica is still scarce and reduced to few summer campaigns. Basille et al. 1997 presented the first studies using radiogenic isotope ratios of ^87^Sr/^86^Sr and ^143^Nd/^144^Nd in insoluble mineral particles in Antarctic ice cores to demonstrate that most dust reaching Central Antarctica derive from the Patagonian semi-desert region. For the Antarctic Peninsula, Dalia et al. (2004) and Pereira et al. (2006) employed simultaneous trace elements in aerosols, ground meteorological databases, acquisition of satellite images and atmospheric transport models to characterize the influence of the South American continent over the Northern Antarctic Peninsula in terms of dust emissions.

Although, the atmospheric dispersion of microorganisms reaching Antarctica have been reported before (Marshall et al. 1996a, 1997), their use as atmospheric tracers, based on quantitative and qualitative diversity in dated ice layers is still scarce. In this work, we have combined the microbiological analysis of bacteria entrapped in snow/firn/ice layers with concomitant tracers of lower latitude continental locations in order to investigate if microorganisms are transported to Antarctica jointly with warmer air masses. For King George Island we employed as tracer the geochemistry of dust material (Ca, Ti and Fe) and for Northern Antarctic Peninsula / Plateau Detroit the stable isotopes of oxygen that exhibit different signatures according to the latitudes.

## 2. MATERIALS AND METHODS

### 2.1. Study sites

Snow and firn samples were collected at 2 sites in Antarctica (Figure 1): (1) in the accumulation zone of an outlet glacier (Wanda Glacier) at Krakow Icefield / King George Island (61°50’-62°15’S; 57°30’-59°00’W) on November 2004, 350 m a.s.l., under the support of the Brazilian Antarctic Program; and at (2) Detroit Plateau located at Graham Land in Northern Antarctic Peninsula at approximately 2,000 m a.s.l. (64°05’S; 59°36’W) during CASA (Climate of the Antarctica and Southern America) project expedition between November 7^th^, 2007 and December 14^th^, 2007 and and ITASE (International Trans Antarctic Scientific Expedition) research initiatives, a joint international scientific collaboration among United States, Chile and Brazil. Both snow pits at the two locations were manually setup by the scientific teams.

**Figure 1.**
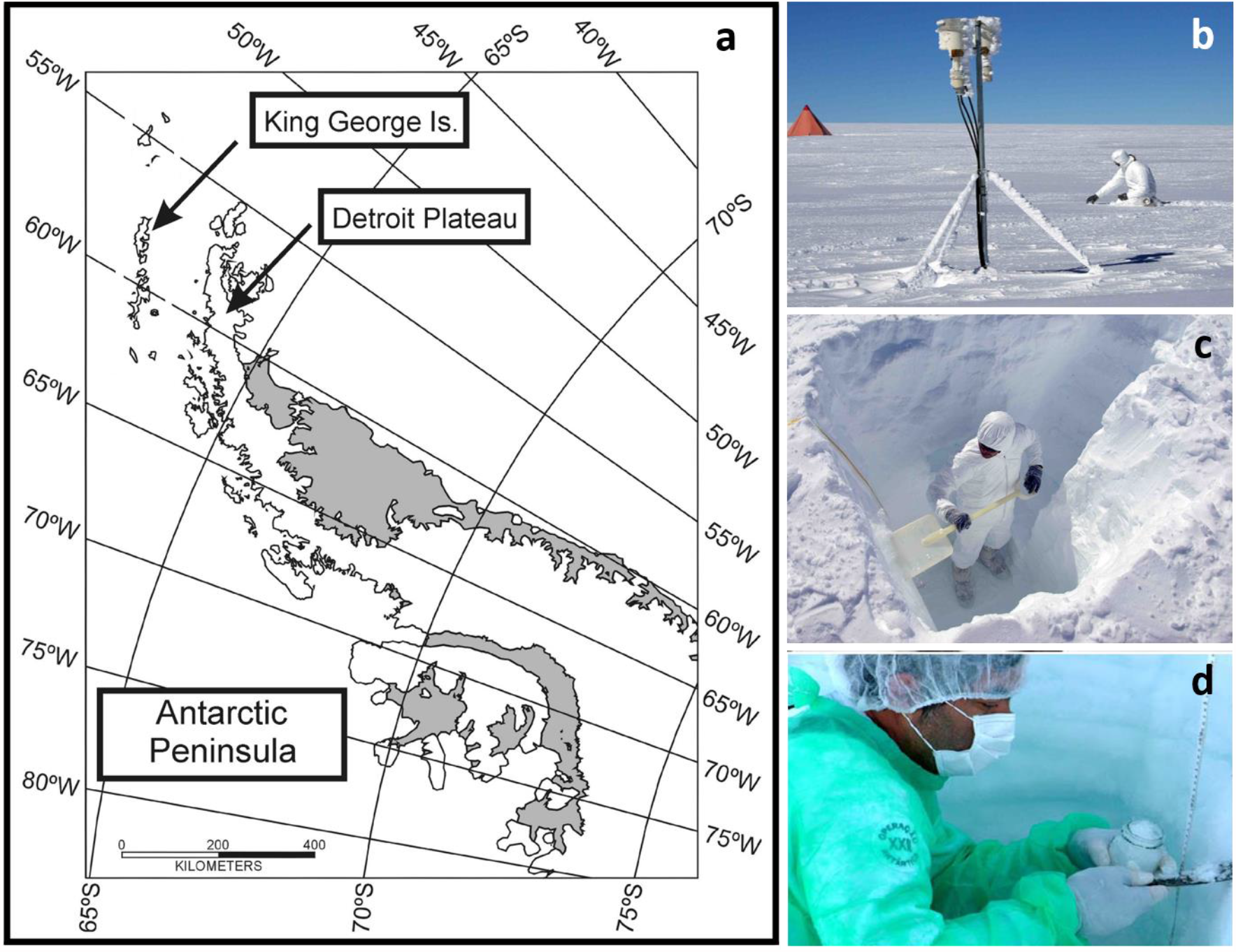
(a) Sampling sites at Antarctic Peninsula for microbiological identification and (b-d) sampling of aerosols and snow/firn layers for microbiology at Detroit Plateau (Northern Antarctic Peninsula) and Wanda Glacier (King George Island).

#### 2.1.1. Samplings at King George Island / South Shetland Islands

The South Shetland Islands/Antarctic Peninsula are located approximately 550 km away from the Southern South America tip and is influenced by climatic regimes of several origins (polar, oceanic and continental, mainly derived from South America), Evangelista (1999). The wind structure is predominantly influenced by the high frequency of cyclonic systems that advect humid and warm air into that region, promoting strong winds and large volume of precipitation (Bintanja, 1995). Evangelista and Pereira (2002) demonstrated that the cyclonic trajectories and their intrinsic energy are the key factors that contribute most to the apportionment of the atmospheric particulates from South America to that site. Along the year mean air temperature ranges from +2oC to - 10°C. At King George Island, summer temperature remains above 0°C, typically from December to the end of March, which is responsible for intense melting processes well evidenced by the ionic and isotopic (δ^18^O) signals in ice cores retrieved from its main domes, Simões et. al (2004). In the present work we sampled in November 2014 before air temperatures reach melting values in King George Island. Therefore, the sampled snow pack corresponded to the period of early autumn when accumulation of snow started in the region to late spring of 2004 and can be considered preserved. Attending the need of retrieving microorganisms from snow/firn samples, three shallow pits (I, II and III) of 140 cm, 200 cm and 80 cm depth were dug on King George Island. Samples were collected on aseptic procedures, Figure 1, storage in autoclaved glass flasks and kept frozen until laboratory procedures in Brazil. Samples were collected at least 1 km away from outcrop rocks, the depth of snow pits were limited by the thick ice layer referred to the summer of the previous year. Samples of Pit I and II were collected in every 20 cm layer while pit III was sampled in each 10 cm. Pits II and III were only used to setup the microbiological method protocols while Pit I consisted of the main sample group which data comprise this work. Mean snow/firn volume for each layer was 150 mL and a total number of 25 samples obtained from the 3 pits.

Accumulation rate of the snow pack was based on the air temperature database obtained at the AWS (Automatic Weather Station) located at the Brazilian Antarctic Station Comandante Ferraz (∼7 km from Wanda glacier). Both are located in the Admiralty Bay in King George Island. The positive to negative temperature turn over, when snow may deposit and start to accumulate, occurred approximately around March 15^th^ which corresponded to the bottom chronology of the snow pack (140 cm from the top). Top of the pits corresponded to the sampling period. An estimate of the months along the snow profile was achieved considering a uniform deposition model. Figure 2 depicts inter-annual data of snow precipitation, wind velocity and air temperature for the study site, during 2004, available at http://www.cptec.inpe.br/antartica/. Instrumental precipitation inferred at the AWS presented an approximate uniform pattern as demonstrated by ANOVA (F=0.31; F_critical_=4.07; P=0.82), calculated at 0.05 significant level for data in the shaded box at Figure 2 (bottom). Small decrease observed during July-August for precipitation may be explained by the increased wind velocity at that period, which is mostly responsible for a drift of surface snow.

**Figure 2.**
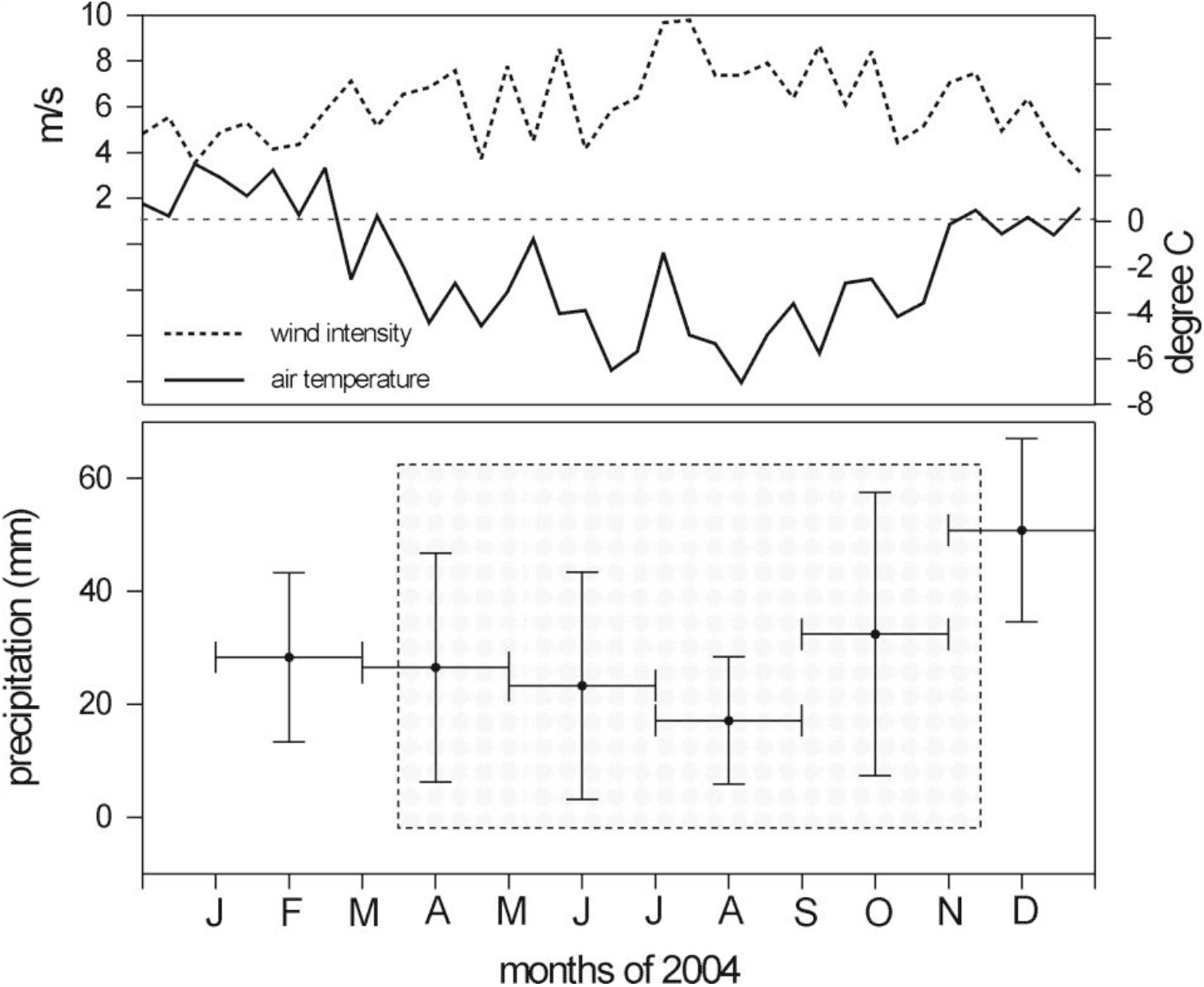
Meteorological data from the AWS at the sampling region. Shaded box bellow represents the period of snow accumulation during 2004.

#### 2.1.2. Detroit Plateau / Northern Antarctic Peninsula

Detroit Plateau is located at Graham Land in Northern Antarctic Peninsula, with heights above 1,500 m. Contrarily to King George Island, very few is known on the climatic and snow accumulation at the plateau. Sampling at Detroit Plateau adopted the same microbiological protocols as those established to Wanda glacier/King George Island. A 2m-pit was dug and snow layers’ chronology was inferred by the δ^18^O measurements in snow replicas. *In situ* meteorological data obtained at Detroit Plateau evidenced that summer temperatures do not reach the melting point.

### 2.2. Cultivation, Isolation and Sequencing of Microorganisms

All cultures were incubated in dark condition, without shaking at 37°C and 16°C for 120 days. Samples were melted at 4°C during a 6 hours period. Aliquots of 20 mL of each sample was taken for culture assays. Two different liquid media were used: Brain Heart Infusion (BHI) and Tioglicolate (TGI). The media were distributed in test tubes (5 mL per tube) and 2x concentrated in order to get the normal concentration after the sample inoculation. To the TGI tubes we added 2.5 mL of mineral oil before sterilization process. A volume of 5 mL of each sample was inoculated in two BHI tubes and two TGI tubes (for different incubation temperatures). After inoculation, BHI cultures were aired in hard shaker procedure for 2 minutes, while in TGI tubes the samples were gently added directly in the media with a pipette thought the oil layer.

Isolate samples were submitted to standard protocol for DNA extraction and purification with the GFX Genomic Blood DNA Purification (Amersham Biosciences). The V3 hypervariable region of the 16S rRNA were amplified and sequenced using the primers 27F (5’AGA GTT TGA TCM TGG CTC AG 3’), (Lane, 1991) and L4101r (GCG TGT GTA CAA GAC CC) (Nudel *et al*., 1996). The PCR profile consisted of 5 min at 95°C, 30 cycles of 1 sec at 95°, 1 min at 60°C, and 2 min at 72°C, with a final extension step for 10 min at 72°C. Sanger sequencing reactions were performed at the MegaBACE 1000 (Amersham Biosciences) platform with the use of the DYEnamic ET Dye Terminator Kit (with the Thermo Sequenase™ II DNA Polimerase), with 25 cycles of 10 sec at 94°C, 5 sec at 50°C and 4 min at 60°C, and the use of a internal primer 338F following Muyzer et al. 1993, 1995). All chromatograms were visually inspected for accuracy and to minimize missing data using the software Geneious v4.82. After the initial screening at the NCBI database using the BLASTn search algorithm, a Neighbor-Joining analysis was conducted with 1.000 bootstrap replicates. The chosen nucleotide substitution model for the dataset was the TN93 based on the Akaike criterion.

### 2.3. Denaturing Gradient Gel Electrophoresis (DGGE)

The PCR products were analyzed by denaturing gradient gel electrophoresis (DGGE) using the standard BioRad protocol with the DCode™ Universal Mutation Detection System. The amplified fragments for the V3 region of the 16S rRNA were analyzed in a vertical polyacrylamide gel (8%) in a 1X TAE buffer, and a sustained 65 °C during the entire electrophoresis.

### 2.4. Flow Cytometry of Snow Samples

Aliquots of 100 mL from each sample were concentrated by freeze drying and re-suspended in 5 mL of PBS 1x pH 7.2. The nucleic acid-specific fluorochrome SYTO 13 green fluorescent nucleic acid stain in 5 mM solution in DMSO (Molecular Probes, Eugene, OR) was used in all experiments. The samples were stained with 20 µM of SYTO 13, vortex to mix then incubate for 1-30 min in the dark at room temperature. The absorption wavelength of SYTO 13 was 488 nm and its emission was 509 nm (Guindulain et al., 1997) All the experiments used a single green fluorescent detector (FL1). Five thousand events were acquired in which sample. The samples were analyzed in an EPICS ALTRA flow cytometer (Beckman Coulter Inc., Hialeah, FL) and analyzed in Expo 32 software (Beckman Coulter).

### 2.5. Elemental Composition Analysis

Aliquots of 400 mL from each 20 cm of Pit I were concentrated by freeze drying process and re-suspended in 10 mL of milliQ™ water followed by filtration in 0.1 μm Nuclepore™ filter. The filtered material was submitted to a Particle Induced X-Ray Emission (PIXE) analysis, owing the concentration of terrigenous elements (Ca, Ti and Fe), previously employed to track continental air masses at the studied site, Correia (1998). In this technique, the filter is exposed to a 2 MeV proton flux bean generated by a Van De Graaff linear particle accelerator, installed at the Institute of Physics of Pontifícia Universidade Católica (PUC-RIO). Details of PIXE technique can be found at Barros Leite et al. 1978. Metal analysis of Ca, Ti and Fe refer to King George Island samples. For Detroit Plateau aerosol samples were collected at daily resolution and analyzed in nuclepore filters (47 mm diameter and 0.4 μm porosity), using an air sampling flux rate of 12 Lpm. Aerosol composition was based on S, Na, Fe and Al elements.

### 2.6. Isotopic analysis

Isotopes in water precipitation have proven to be a tracer in studies of the interaction between air masses and synoptic climatology, especially in terms of origin of the water vapor. The spatial distribution of global δ^18^O, as provided by the Global Network of Isotopes in Precipitation (GNIP)/IAEA (accessible at *http://isohis.iaea.org*), show an isotopic signature of d^18^O ranging from -15 ‰ to -3.0 ‰ for the tropical to temperate sites of the South Hemisphere and values corresponding to -15 ‰ to -30 ‰ for polar air masses. Therefore δ^18^O in snow may tag the presence of warmer or colder incursions of air masses at the Maritime Antarctica where our work was developed.

The device used for δ^18^O analysis was an automatic interface of the DeltaPlus Advantage gas source mass spectrometer of the Thermo Finningan™ brand. A fraction of each liquid sample, referring to a 10 cm snow layer was processed for the analysis. The international standards used were: V-SMOW, IAEA-SLAP2 and IAEA-GISP2. δ^18^O analysis refer to Detroit Plateau samples only.

### 2.7. Atmospheric transport model

In order to track the air mass past migrations towards Antarctica we have used the HYSPLIT (Hybrid Single Particle Lagrangian Integrated Trajectory) model. The HYSPLIT model is an atmospheric trajectory modeling tool widely used to help understand atmospheric transport, dispersion and deposition of mineral dust and pollutants. HYSPLIT is a platform developed by ARL (Air Resources Laboratory), belonging to NOAA (National Oceanic and Atmospheric Administration) and available online (http://ready.arl.noaa.gov/HYSPLIT.php).

The method of calculating the model is a hybrid between the Lagrange approach (using a mobile reference for advection and diffusion calculations as the trajectories move from their initial location) and the Eulerian methodology (which uses a fixed three-dimensional grid as a frame reference points for calculating pollutant concentrations in the air). The model output depicts trajectories of the center of the air masses backwards in time and allow inferring the air mass transit before reaching the study site.

## 3. RESULTS AND DISCUSSION

### 3.1. Identification of microorganisms for King George Island

From 20 inoculated snow samples from King George Island, bacterial growth was observed in 10 (3 only on BHI incubated at 37 °C, 1 only on BHI at 16 °C, 2 only on TGI at 16 °C, 1 on BHI and TGI 16 °C, 2 on BHI 16 °C and 37 °C; 1 on BHI and TGI 16 °C and 37 °C. All cultivated microorganisms were identified as gram-positive bacteria, and more than 80% of those were spore-forming *Bacillus*. They were isolated and submitted to molecular sequencing techniques which results are presented in Figure 3 and Table 1.

**Figure 3.**
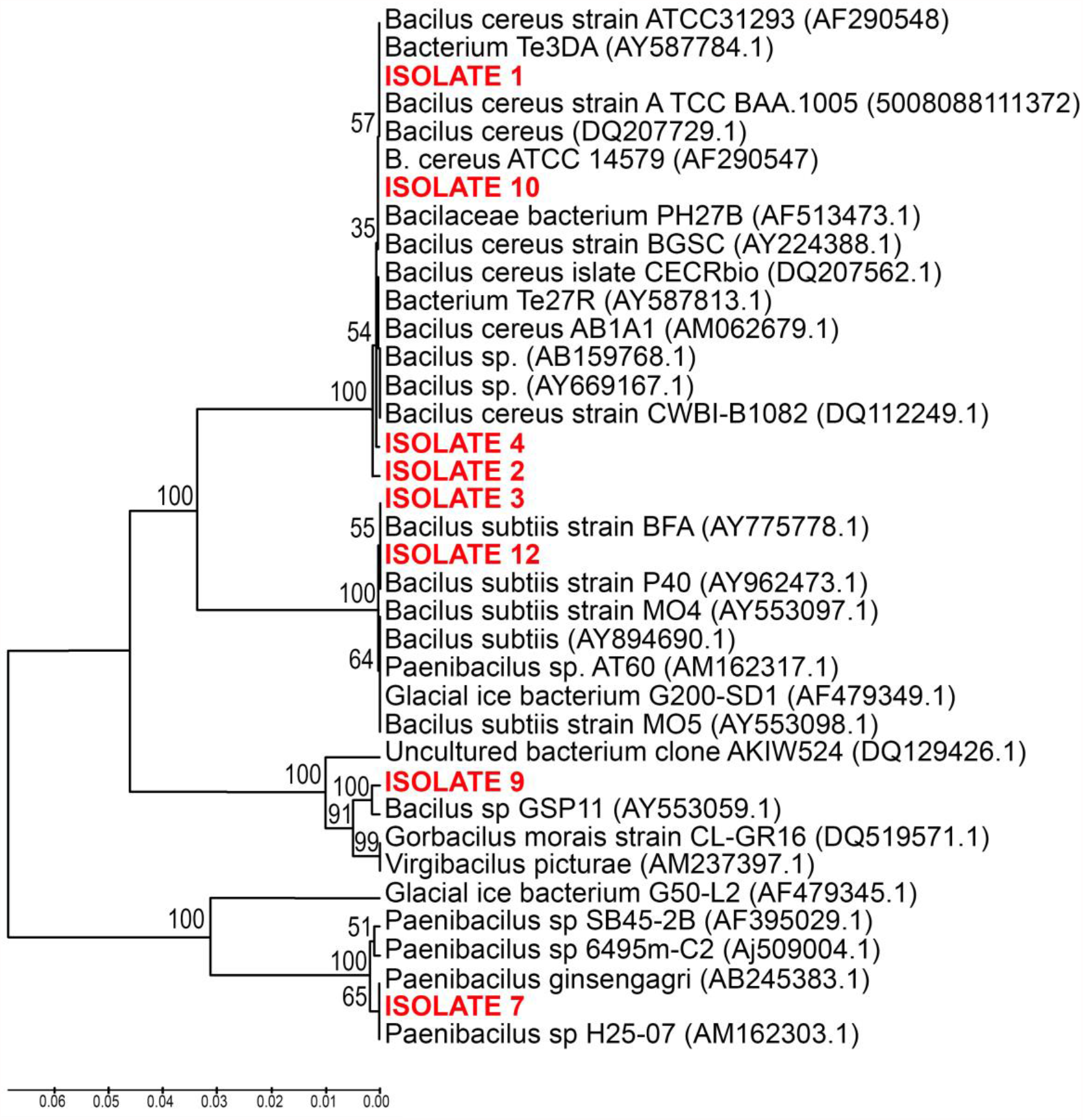
Neighbor-joining 16S rRNA tree (TN93) (bootstrap x1000) of the isolated bacilli, marked in red.

**Table 1.** Genetic identification of the isolated bacilli. The table presents the name of the isolate, the size in base pairs of the analyzed sequence, the GenBank classification for the sequence and its homology within the previous published strains.

| Isolate ID | Size (BP) | GenBank | Homology (%) |
| --- | --- | --- | --- |
| 1 | 1310 | *Bacillus cereus* | 1304/1304 (100%) |
| 2 | 503 | *Bacillus* sp. | 473/485 (97.5%) |
| 3 | 1284 | *Bacillus subtilis* | 1278/1278 (100%) |
| 4 | 874 | *Bacillus cereus* | 872/874 (99%) |
| 7 | 1282 | *Paenibacillus* sp. | 1282/1285 (99.7%) |
| 9 | 939 | *Bacillus* sp. | 934/939 (99.4%) |
| 10 | 1289 | *Bacillus cereus* | 1289/1289 (100%) |
| 12 | 1281 | *Bacillus subtilis* | 1281/1281 (100%) |

The resultant Neighbor-joining tree (Figure 3) show the isolates 1, 10, 4,and 3 recovered nested within the *Bacillus cereus* group, isolates 3 and 12 within the *Bacillus subtilis* group, the isolate 7 recovered nested within the *Paenibacillus* group, and the isolate 9, recovered in a group together with *Goribacillus litoralis* and *Virgibacillus picturae*.

Among the sequenced isolates, two of them were identified to the species level (*Bacillus*) and the remaining to the genus level (*Bacillus* and *Paenibacillus*). Among the *Bacillus* isolates, *Bacillus cereus* group if formed by about 11 closely related species with contributions for production of numerous enzymes (Chang et al. 2007), metabolites (Kevani et al. 2009), removal of various heavy metals (Chen et al. 2016), persistent organic pollutants (Kazunga and Aitken, 2000), and as probiotics for humans and animals (Gisbert et al. 2013). The *B. cereus* bacteria group occupy several habitats ranging from terrestrial and aquatic environments, and also including plants and animals, and presents strong survivability of spores allowing them to better withstand hostile conditions and to better disperse (Guinebretiere et al. 2008, Jensen et al. 2003).

The second isolated and identified *Bacillus* species, *B. subtilis* can be isolated from many environments such as terrestrial and aquatic. The species is ubiquitous and broadly adapted to grow in diverse settings within the biosphere. In response to nutrient deprivation, *B. subtilis* can form highly resistant dormant endospores (Sonenshein et al. 2002, Ricca et al. 2004). These spores could easily disperse by wind (Merrill et al. 2006, Jaenicke (2005) and migrate long distances. *B. subtilis* is also referred to as a soil dweller, likely presenting a saprophytic life history (Vilain et al. (2006). *B. subtilis* can also grow in close association with plant roots, and also within the gastrointestinal tract of several animals (Tam et al. 2006, Leser et al. 2008, Hong et al. 2005, Inatsu et al. 2006). In marine environments, although growth might occur, the abundance seems to be related with its observed association with the gastrointestinal tract of marine organisms (Newaj-Fyzul et al. 2007) and other biotic surfaces (Ivanova et al. 1999). Current results indicate that the apparent ubiquity of *B. subtilis* seems to be not only a result of spore persistence in these environments since *B. subtilis* seems to actively grow in diverse environments including ranging from soils, water, gastrointestinal tracts, plant roots.

The third identified isolate is related with *Paenibacillus*. The genus *Paenibacillus* (Ash et al. 1993) was created to receive the former ‘group 3’ species of the genus *Bacillus*. It includes about 35 species of facultative anaerobes and endospore-forming, neutrophilic, periflagellated heterotrophic, low G+C gram-positive bacilli. *Paenibacillus* species were recorded from different environments such as soils, roots, rhizosphere of various crop plants including wheat, sorghum, sugarcane, maize, barley (Guemouri-Athmani et al. 2000, von der Weid et al. 2000), trees such as pines (Holl and Chanway, 1992, Shishido et al. 1996), and also marine sediments (Ravi et al. 2007).

Yet those bacteria, such as *B. subtilis* or *B. cereus*, can be found in many different substrata and conditions, they do not have specific mechanisms to allow them to maintain a normal metabolism and cell division with respect to the relative low temperatures as those of the polar environments. The presence of those non-natural cold environmental flora microorganisms, and the long time in incubation (up to 60 days in some cases), contrasting with 24 hours of growth, after the re-inoculation, suggests its exogenous origin. After a long period deposited on ice, the cells and the spores remained viable but far from its optimum metabolic conditions, explaining their long time to recover in incubation.

### 3.2. Microorganism Counting, DGGE, and Elemental analysis for King George Island

Counting rate obtained by the flow cytometry showed a variable cell distribution with a pronounced peak at the 60-80 cm layer and two secondary peaks, at 20-40 cm and 120-140 cm, Figure 4. The major peak being four-fold higher than the remaining data. The concentration of unmarked particles and DNA marked cells had a similar variation for the analyzed layers. Of special interest was the data provided by the PIXE analysis which showed an impressive increase of insoluble inorganic elements (Fe, Ti and Ca) between 60-100 cm snow layers, and therefore enclosing the peak of microorganisms. This nearly concomitant peak strongly suggests that mineral particles and microorganisms were transported in the same synoptic event to the Antarctic region. Such association between alfresco bacterial communities and resuspended soil dust was previously reported and detailed in previous studied (Savoie 1989; Li et al. 1996; Prospero 1996, 2003; Griffin et al. 2001, 2003). Lighthart and Stetzenbach (1994) reported that bacteria and fungi can be transported by wind for long distances and remaining viable over normal environmental conditionings. Lighthart and Shaffer (1997) pointed out to the importance of aerosol droplet/particle in the maintenance of cell viability along an atmospheric transport.

**Figure 4.**
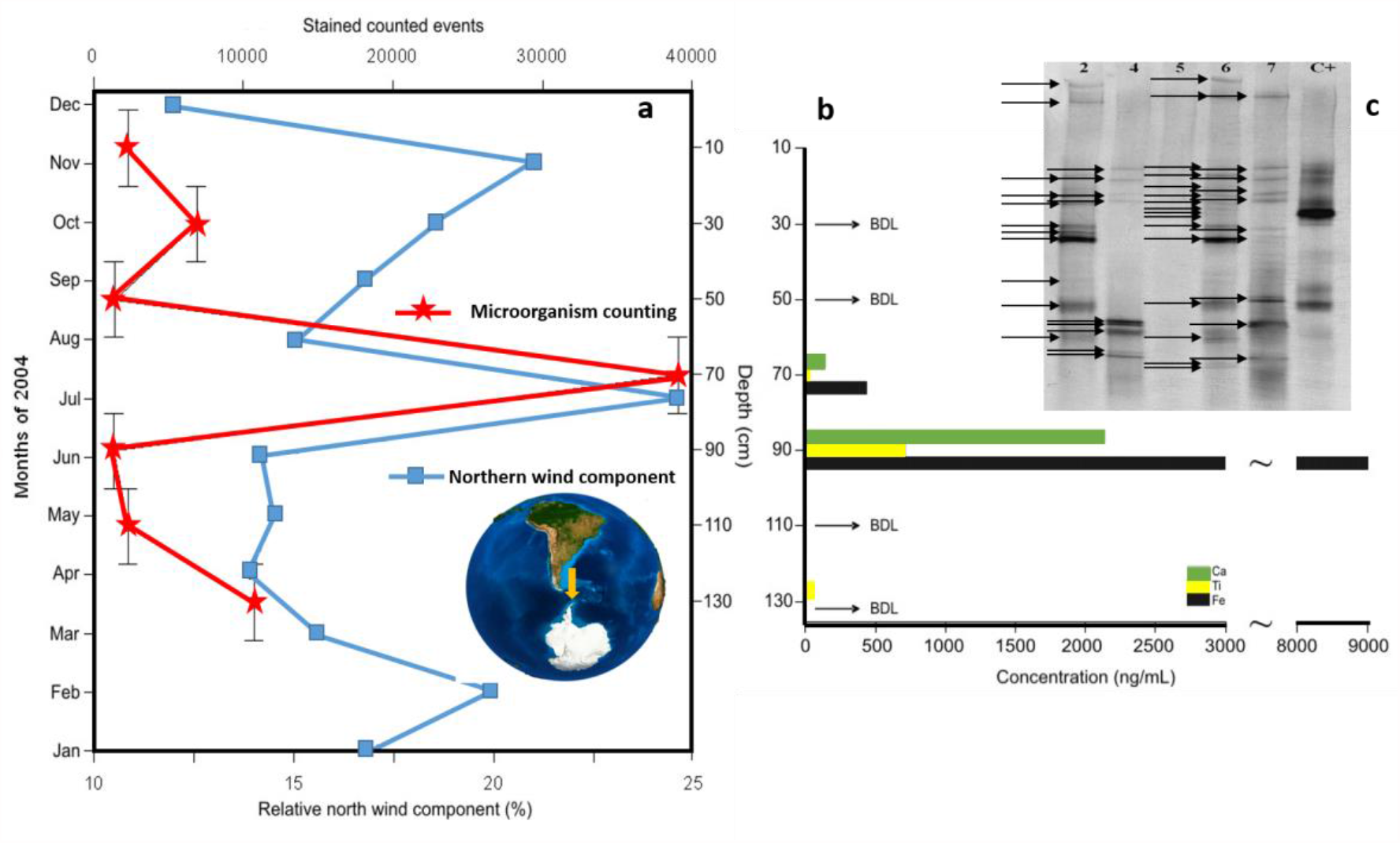
Microorganism counting from a shallow snow pit in Wanda Glacier (a) and relative north wind component at King George Island (according to snow deposition chronology); crustal elements Ca, Ti, Fe concentrations at same samples (b) (BDL: bellow detection limit); DGGE polyacrylamide gel (8%) obtained from the pit I at Wanda Glacier. The lanes correspond to sections: (2) 20-40 cm – 11 bands; (3) 60-80 cm -9 bands; (4) 80-100 cm – not detected; (5) 100-120 cm – 15 bands; (6) 120-140 cm – 10 bands; C+ (positive control). Each observed band is marked by a black arrow.

As an attempt to investigate the atmospheric event associated with the peaks of microorganisms and crustal elements, we combined a set of information: *in situ* meteorological data (wind direction), satellite image galleries around latitude 60oS and the air masses back-trajectories from the HYSPLIT model. A detailed study of Evangelista (1999) concerning individual atmospheric transport events between South America and the Antarctic Peninsula stressed the high association between the meridional crustal material transport towards Antarctica and the northern wind component. On basis of this previous result, we have overlaid microorganism counting, the north wind component and crustal elemental concentrations assuming a uniform snow deposition chronology. The result shows a consistent association among the microbiological, meteorological and geochemical parameters.

According to the snow pit chronology, the peak of microorganisms occurs at middle winter, which confirms the hypothesis of atmospheric transport and deposition, during that season contamination due to weathering of ice-free areas by surface winds, presence of polar animals, and human occupation get their minimum values. Additionally, our DGGE results also suggest high diversity on layers from 100-140 cm. The strata correspond to March-April period, at the end of the Antarctic summer of 2004, when the Antarctic ice cap is not entirely formed and the resuspension of the local microbiota is expected due to the strong and constant surface winds.

Within this period, we have selected 215 satellite images from GOES satellite in ftp://ftpantartica.cptec.inpe.br/pub/antartica/GOES_SubA/ at the Infra-Red and visible bands and we have run the HYSPLIT back-trajectories model for each day. From the image gallery and the atmospheric transport model, we have recognized events during July 2014 in which air mass trajectories previously have passed through the South American continent before reaching King George Island. One may observe that this period is coincident with the increased north wind component observed in the Meteorological Station at King George Island, Figure 4. In that circumstance, the air mass could be enriched by dust and microorganisms during their continental migration. From this method we have selected three potential events attending the above characteristics. Figure 5 depicts simultaneous GOES images and the HYSPLIT back-trajectories model output for three advective episodes linking South America and Antarctica Peninsula.

**Figure 5.**
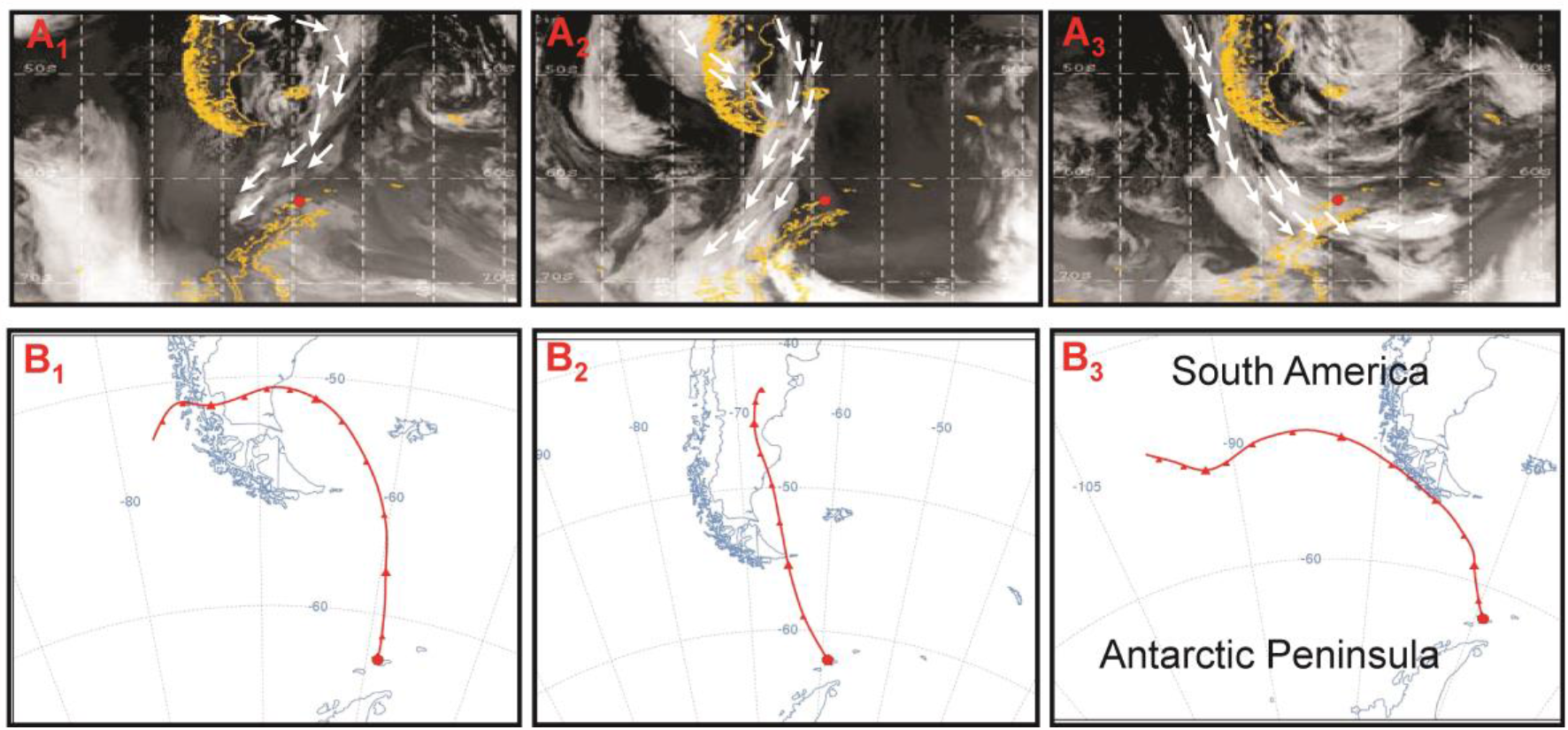
Satellite images and the HYSPLIT back-trajectories for 3 southwards air mass advection episodes between South America and Antarctic Peninsula. Episode (A1,B1) occurred in 8^th^ July, 2004; (A2,B2) in 19^th^ July, 2004; (A3,B3) in 22^nd^ July, 2004. Small red dot is the location of King George Island. The superimposed arrows were defined on basis of corresponding synoptic charts analysis for the same days.

From Figures 4 and 5, it is clear the correspondence among the parameters used to track the atmospheric transport. The three events of southwards air mass advection originated in the meridional Patagonia semi-desert sector, may explain consistently the elevated concentrations of crustal elements found in the snow samples. Dust plume spread over the Southern Ocean has been detailed described from satellite measurements of MODIS and OMI/NASA indicating the large influence of Patagonia to the delivery of dust to Sub-polar sites (Gasso and Stein 2007) and its relevance to the ocean primary productivity through Fe deposition (Erickson et al., 2003). Although limited to one sequence of episodes of transport, this work provides an initial basis for further investigation in deeper ice core of that region, which could bring additional knowledge on microorganisms apportioning polar regions.

### 3.3. Microorganism counting and Isotope analysis for Detroit Plateau

During the Detroit Plateau mission, we collected aerosols *in situ* in order to characterize their behavior at high altitudes and its relation with the atmospheric transport. We sampled during 6 different days integrated for 24 hours. Elemental composition was based on Na and S as representative of marine influence and Fe and Al as representative of crustal mineral material. The results present in Figure 6 show that concentration of crustal material is closely related to the nature of the air mass. The episodes of 3, 4 and 5^th^ December 2007 presented significantly higher concentrations of Al and Fe coincident to air masses which migration history is associated to the Southern Patagonia, evidencing a similar mechanism as observed to King George Island.

**Figure 6.**
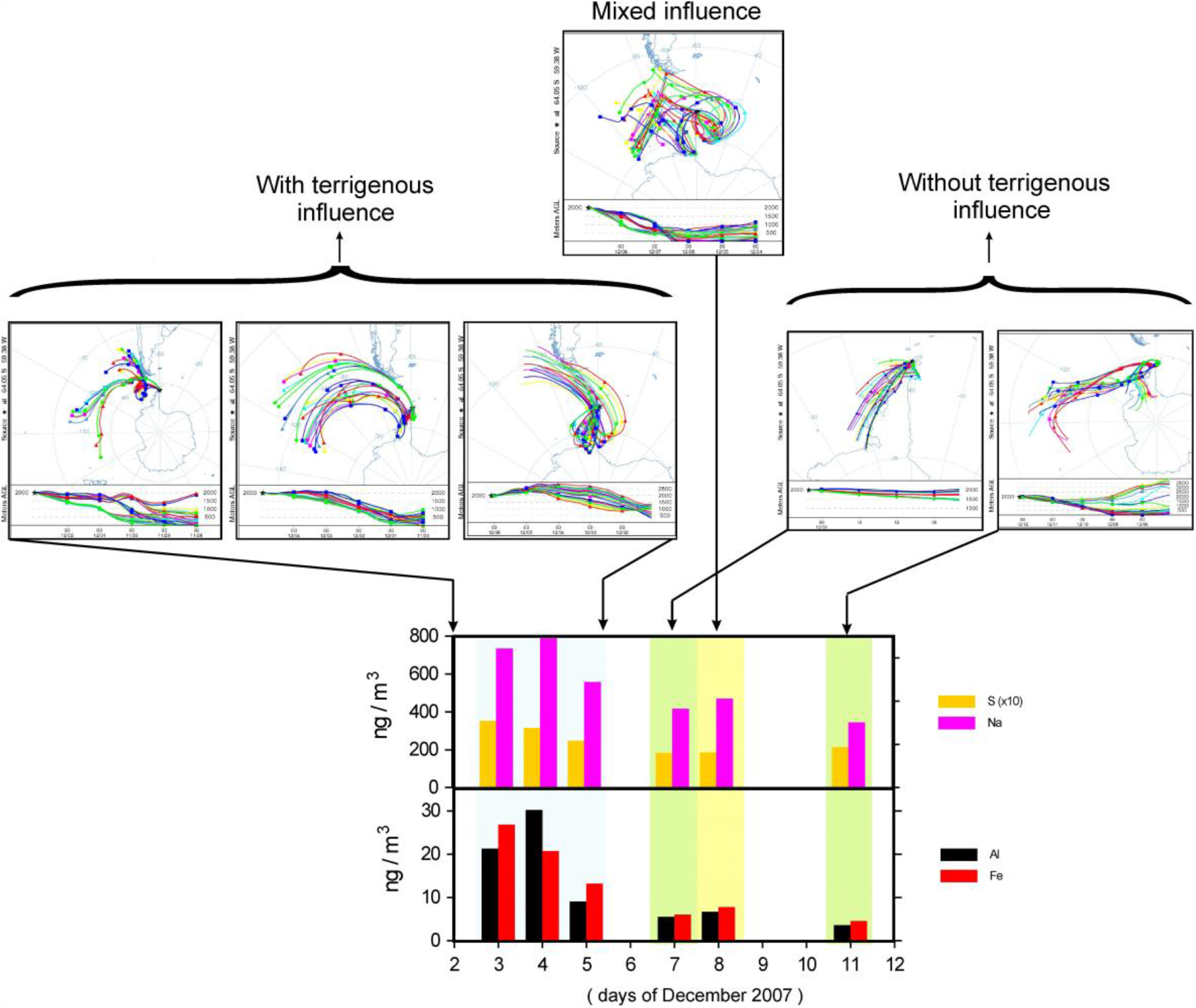
Elemental composition for aerosols at Detroit Plateau during the 2007 campaign and associated air mass backward trajectories. “Terrigenous influence” refers to the migration of the air mass over continental land of South America.

With respect the Detroit plateau 2m-snow pit, microorganism counts and corresponding oxygen isotopic data covariate well and corroborated the concept of atmospheric transport. Stable isotopes of oxygen, δ^18^O, in precipitation is influenced by a combination of factors, such as precipitation amount, air temperature, moisture source, altitude and vapor transportation. The northern Antarctic Peninsula is particularly complex in the meteorological and synoptical point of view, since it is a place where exist multiple influences of different air masses (Polar, Southern Pacific and Southern American). Data of δ^18^O in the snow pit of Detroit plateau varied from -14‰ to -27‰. Microorganism counts varied in a very close pattern as δ^18^O, Figure 7. Increases of microorganism counts occurred in layers where δ^18^O values were representative of lower latitudes in the Southern South America while values bellow -20 ‰ reflect the polar influence in the region. Interesting to observe is that when the polar influence is present, microorganism counts is reduced by a factor between 2.5 and 3.0 probably indicating that the (sub)polar environment can be also a source of microorganism and constitute a background level at that region.

**Figure 7.**
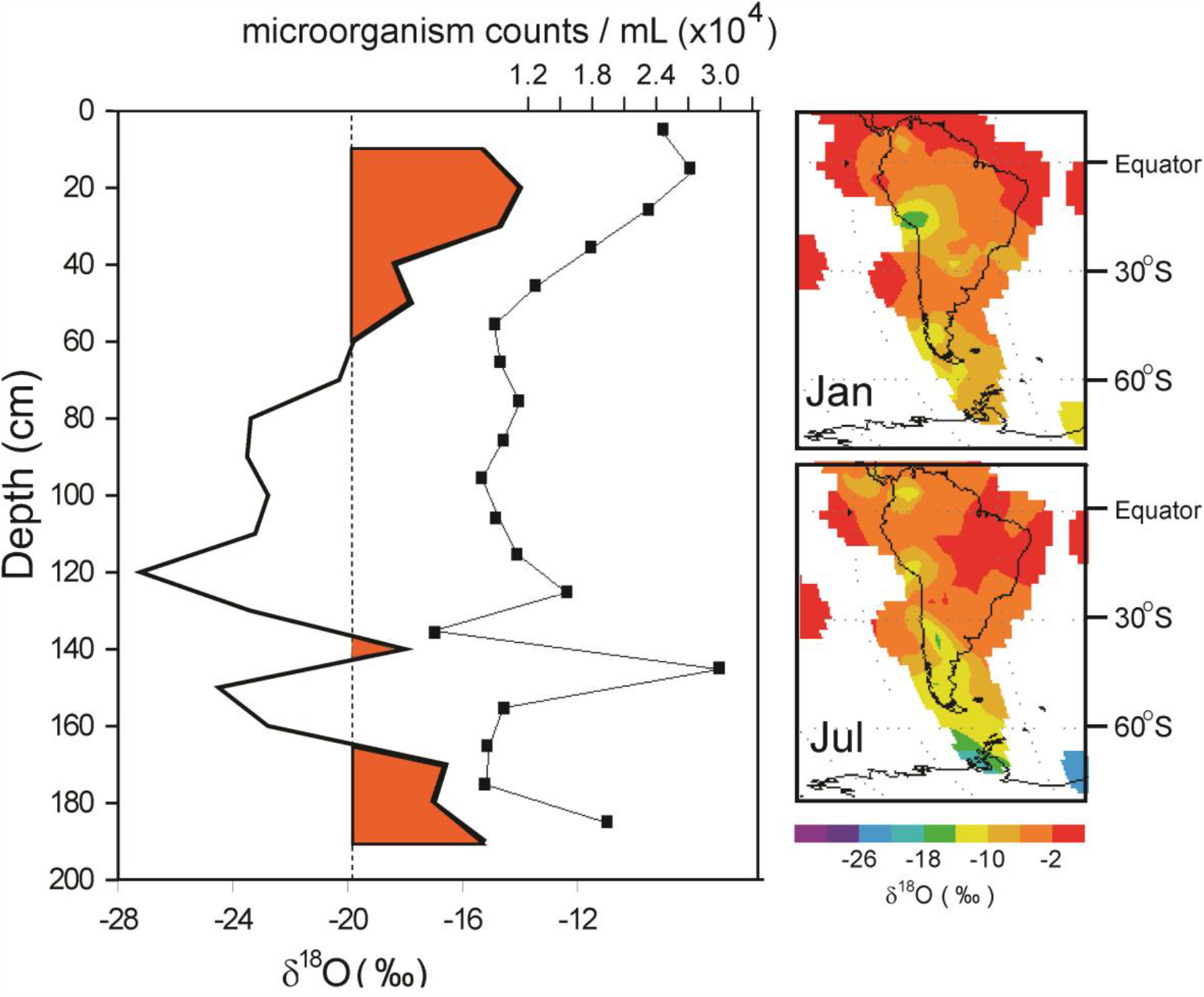
(left) Microorganism counts and δ^18^O. Colored parts of δ^18^O curve indicates possible warm air advection from lower latitudes; (right) average δ^18^O in precipitation from AIEA/GNIP for January (summer) and July (winter) periods.

## 4. CONCLUSIONS

In this work we explored the use of molecular identification and counting in snow/firn layers of Antarctic ice sheets, by flow cytometry, that is a simple and technological facility well available in several laboratories around the world. We developed simultaneous measurements of microorganisms and other geochemical tracers successfully used in the literature as the elemental composition, representing tracers of crustal material based on Fe, Ti, Al and Ca, and stable isotopes of oxygen. We found satisfactory associations between microorganisms and the geochemical tracers that were confirmed by atmospheric dispersion models and satellite images. We conclude that same events that transport dust to Antarctica are source of microorganisms that are probably associated to dust resuspension in the continents around Antarctica, especially South America. Therefore, microorganism counting in ice sheet layers can act as tracer of the continental terrigenous influence, their desertification evolution and wind dynamics between the continents.

## ACKNOWLEDGMENTS

We thank Ivan and Denilson for helping with the sterilization of the sampling equipment. We thank FAPERJ to have provided scholarship. This research was funded by CNPq/PROANTAR (process: 55.0353/2002-0) and logistics provided by SCIRM and CAP (Clube Alpino Paulista). We also thank the CASA project and University of Maine to have provided the samplings at Detroit Plateau and measurements of the stable isotopes of oxygen.

## AUTHOR CONTRIBUTIONS

**MC** – Investigation, Writing – original draft

**HE** – Conceptualization, Supervision, Funding acquisition, Project administration

**JAAP** – Resources

**ALB** – Resources

**VP** – Supervision, Resources

**EK** – Investigation

**MS** – Investigation, Development of the snow samplers and field sampling at Wanda Glacier, Antarctica

**KDC** – Investigation

**ASA** – Investigation

**CA** – Investigation, Writing – original draft, Writing – review & editing

## REFERENCES

Abyzov SS. 1993. Microorganisms in the Antarctic ice, In: E I Friedman (ed) Antarctic microbiology, Wiley-Liss, New York, pp 265–295

Ash C, Priest FG and Collins MD. 1993. Molecular identification of rRNA group 3 bacilli (Ash, Farrow, Wallbanks and Collins) using a PCR probe test. Proposal for the creation of a new genus Paenibacillus. Antonie Van Leeuwenhoek 64:253–260

Basille I, Grousset FE, Revel R et al. 1997. Patagonian origin of glacial dust deposited in East Antarctica (Vostok and Dome C) during glacial stages 2, 4 and 6. Earth and Planetary Science Letters 147:573–589

Bidle KD, Lee S, Marchant DR et al. 2007. Fossil genes and microbes in the oldest ice on Earth. Proceedings of the National Academy of Sciences of the United States of America 104:(33) 13455–13460

Bintanja R. 1995. The local energy surface balance of the Ecology Glacier, King George Island, Antarctica, measurements and modeling, In: Bintanja R (ed) The Antarctic ice sheet and climate, Amsterdam, Utrecht University, pp 41–59

Chang WT, Chen YC & Jao CL. 2007. Antifungal activity and enhancement of plant growth by Bacillus cereus grown on shellfish chitin wastes. Bioresour Technol 98, 1224–1230.

Chen, Z., Pan, X., Chen, H., Guan, X. & Lin, Z. 2016. Biomineralization of Pb(II) into Pb-hydroxyapatite induced by Bacillus cereus 12-2 isolated from Lead-Zinc mine tailings. J Hazard Mater 301, 531–537.

Christner B, Mosley-Thompson E, Thompson LG et al. 2003. Bacterial recovery from ancient glacial ice. Environmental Microbiology 5(5):433–436

Dalia KC, Evangelista H, Simões J et al. 2004. Sazonalidade de aerossóis atmosféricos e microanálise individual por EDS em testemunho de gelo da ilha Rei George. Brazilian Antarctic Research 4:25–36

Evangelista H. 1999. Um estudo sobre o transporte atmosférico na ilha Rei George. Tese de Doutoramento. Rio de Janeiro, Universidade Estadual do Estado do Rio de Janeiro. P. 198.

Evangelista H & Pereira EB. 2002. Radon Flux at King George Island, Antarctic Peninsula. Journal of Environmental Radioactivity 61:283–304

Erickson DJ, Hernandez JL, Ginoux P, Gregg WW, Mcclain C & Christian J. 2003. Atmospheric iron delivery and surface ocean biological activity in the Southern Ocean and Patagonian region. Geophysical Research Letters 30 (12), 1609, doi:10.1029/2003GL017241.

Fierer N. 2008 Microbial biogeography: patterns in microbial diversity across space and time. In: Accessing Uncultivated Microorganisms: From the Environment to Organisms and Genomes and Back, ed. K. Zengler (Washington DC: ASM Press).

Gasso S, Stein AF. 2007. Does dust from Patagonia reach the sub-Antarctic Atlantic Ocean? Geophysical Research Letters 34, L01801, doi:10.1029/2006GL027693.

Gisbert E, Castillo M, Skalli A, Andree KB & Badiola I. 2013. Bacillus cereus var. toyoi promotes growth, affects the histological organization and microbiota of the intestinal mucosa in rainbow trout fingerlings. J Anim Sci 91, 2766–2774.

Griffin DW, Garrison VH, Herman JR et al. 2001. African desert dust in the Caribbean atmosphere: Microbiology and public health. Aerobiologia 17:203–213

Griffin DW, Kellogg CA, Garrison VH et al. 2003. Atmospheric microbiology in the northern Caribbean during African dust events. Aerobiologia 19:143–157

Guemouri-Athmani S, Berge O, Bourrain M, Mavingui P, Thiery JM, Bhatnagar T & Heulin T. 2000. Diversity of Paenibacillus polymyxa in the rhizosphere of wheat (Triticum durum) in Algerian soils. Eur J Soil Biol 36:149–159

Guindulain T, Comas J, Vives-Rego J. 1997. Use of Nucleic Acid Dyes SYTO-13, TOTO-1, and YOYO-1 in the Study of Escherichia coli and Marine Prokaryotic Populations by Flow Cytometry. Applied and Enviromental Microbiology 63:(11) 4608–4611

Guinebretiere MH et al. 2008. Ecological diversification in the Bacillus cereus Group. Environ Microbiol 10, 851–865.

Holl FB & Chanway CP. 1992. Rhizosphere colonization and seedling growth promotion of lodgepole pine by Bacillus polymyxa. Can J Microbiol 38:303–308

Hong HA et al. 2005. The use of bacterial spore formers as probiotics. FEMS Microbiol. Rev. 29, 813–835

International Atomic Energy Agency. 2001. GNIP Maps and Animations, International Atomic Energy Agency, Vienna. Accessible at http://isohis.iaea.org

Inatsu Y et al. 2006. Characterization 22 HONG, H.A. AND DUCLE, H. 2004. The fate of ingested spores. In Bacterial Spore Formers: Probiotics and Emerging Applications (Ricca E et al., eds), pp. 107–112, Horizon Scientific Press

Ivanova EP et al. 1999. Characterization of Bacillus strains of marine origin. Int. Microbiol. 2, 267–271

Jaenicke R. 2005. Abundance of cellular material and proteins in the atmosphere. Science 308, 73.

Jensen GB, Hansen BM, Eilenberg J & Mahillon J. 2003. The hidden lifestyles of Bacillus cereus and relatives. Environ Microbiol 5, 631–640.

Karl DM, Bird DF, Bjorkman K et al. 1999. Microorganisms in the accreted ice of Lake Vostok, Antarctica. Science 286:2144–2147

Kazunga C & Aitken MD. 2000. Products from the incomplete metabolism of pyrene by polycyclic aromatic hydrocarbon-degrading bacteria. Appl Environ Microbiol 66, 1917–1922.

Kevany BM, Rasko DA & Thomas MG. 2009. Characterization of the complete zwittermicin A biosynthesis gene cluster from Bacillus cereus. Appl Environ Microbiol 75, 1144–1155.

Lane DJ. 1991: In Nucleic Acid Techniques in Bacterial Systematics E. Stackebrandt, M. Goodfellow, Eds. (Wiley & Sons, Chichester) pp. 115–175.

Leser TD et al. 2008. Germination and outgrowth of Bacillus subtilis and Bacillus licheniformis spores in the gastrointestinal tract of pigs. J. Appl. Microbiol. 104, 1025–1033

Li X, Maring H, Savoie D et al. 1996. Dominance of mineral dust in aerosol light scattering in the North Atlantic trade winds. Nature 380:416–419

Lighthart B & Stetzenbach LD. 1994. Distribution of microbial bioaerosol. In: Lighthart B, Mohr AJ (ed) Atmospheric Microbial Aerosols, Theory and Applications, Kluwer Academic Publishers, Boston, pp 68–98

Lighthart B, Shaffer BT. 1997. Increased airborne bacterial survival as a function of particle content and size. Aerosol Science and Technology 27:(3) 439–446

Marshall A. 1996a. Biological particles over Antarctica. Nature 383:680

Marshall A. 1997. Seasonality in Antarctic airborne fungal spores. Applied and Environmental Microbiology 63:2240–2245

Merrill L et al. 2006. Composition of Bacillus species in aerosols from 11 U.S. cities. J. Forensic Sci. 51, 559–565

Mishra VK, Kim KH, Hong S et al. 2004. Aerosol composition and its sources at the King Sejong Station, Antarctic peninsula. Atmospheric Environment 38:4069–4084

Muyzer G, De Waal EC & Uitterlinden AG. 1993. Profiling of complex microbial populations by denaturing gradient gel electrophoresis analysis of polymerase chain reaction amplified genes coding for 16S rRNA. Appl Environ Microbiol. vol. 59, p. 695–700.

Muyzer G, Teske A & Wirsen CO. 1995. Phylogenetic relationships of Thiomicrospira species and their identification in deep-sea hydrothermal vent samples by denaturing gradient gel electrophoresis of 16S rDNA fragments. Arch Microbiol. vol. 164, p. 165–172.

Newaj-Fyzul A et al. 2007. Bacillus subtilis AB1 controls Aeromonas infection in rainbow trout (Oncorhynchus mykiss, Walbaum). J. Appl. Microbiol. 103, 1699–1706

Nudel U, Engelen E & Felske A. 1996. Sequence Heterogeneities of Genes Encoding 16S rRNAs in Paenibacillus polymyxa Detected by Temperature Gradient Gel Electrophoresis. Journal of Bacteriology. vol. 178, p. 5636–5643.

Pearce DA, Hughes KA, Lachlan-Cope T, Harangozo SA, Jones AE. 2010. Biodiversity of airborne microorganisms at Halley Station, Antarctica. Extremophiles 14, 145–159. doi:10.1007/s00792-009-0293-8

Pearce DA, Alekhina IA, Terauds A, Wilmotte A, Quesada A, et al. 2016. Aerobiology Over Antarctica – A New Initiative for Atmospheric Ecology. Front. Microbiol. 7:16. doi: 10.3389/fmicb.2016.00016

Pereira EB. 1999. Radon-222 Time series measurements in the Antarctic Peninsula (1986-1987). Tellus 42b:(1)39–45

Pereira EB, Evangelista H, Pereira KCD et al. 2006. Apportionment of black carbon in the South Shetland Islands, Antarctic Peninsula. Journal of Geophysical Research 111:(D3)

Priscu, JC, Fritsen CH, Adams EE et al. 1998. Perennial Antarctic Iake ice: An oasis for life in a polar desert. Science 280:2095–2098

Prospero JM. 1996. The atmospheric transport of particles to the ocean. In: Ittekkott V, Honjo S, Depetris PJ (ed) Particle Flux in the Ocean. SCOPE Report 57. John Wiley, New York, pp 19–52

Prospero JM, Lamb PJ. 2003. African droughts and dust transport to the Caribbean, climate change implications. Science 302:1024–1027

Prospero JM. 2005. Interhemispheric transport of viable fungi and bacteria from Africa to the Caribbean with soil dust. Aerobiologia 21:1–19

Ravi AV, Musthafa KS, Jegathammbal G, Kathiresan K and Pandian SK. 2007. Screening and evaluation of probiotics as a biocontrol agent against pathogenic Vibrios in marine aquaculture. Lett Appl Microbiol 45:219–223

Ricca E et al. eds. 2004. Bacterial Spore Formers: Probiotics and Emerging Applications, Horizon Scientific Press

Savoie DL, Prospero JM, Saltzman ES. 1989. Non-seasalt sulfate and nitrate in tradewind aerosols at Barbados, evidence for long-range transport. Journal of Geophysical Research 94:5069–5080

Shishido M, Massicotte HB and Chanway CP. 1996. Effect of plant growth promoting Bacillus strains on pine and spruce seedling growth and mycorrhizal infection. Ann Bot.77:433–441

Simões JC, Ferron FA, Bernardo RT. 2004. Ice core study from the King George Island, South Shetland, Antarctica. Brazilian Antarctic Research 4:9–23

>Sonenshein AL et al. eds. 2002. Bacillus subtilis and Its Closest Relatives: From Genes to Cells, ASM Press

Tam NK et al. 2006. The intestinal life cycle of Bacillus subtilis and close relatives. J. Bacteriol. 188, 2692–2700

Vilain S et al. 2006. Analysis of the life cycle of the soil saprophyte Bacillus cereus in liquid soil extract and in soil. Appl. Environ. Microbiol. 72, 4970–4977

Von Der Weid IA, Paiva E, Nobrega A, Van Elsas JD & Seldin L. 2000. Diversity of Paenibacillus polymyxa strains isolated from the rhizosphere of maize planted in Cerrado soil. Res Microbiol 151:369–381

Warwick FV. 1988. Microbial ecosystems of Antarctica. Cambridge University Press, 303 pp

Warwick FV. 2000. Evolutionary origins of Antarctic microbiota, invasion, selection and endemism. Antarctic Science 12:(3) 374–385.

